# Self-organized vascularized human liver spheroids: Serum-free culture conditions and use as tissue building blocks

**DOI:** 10.1101/2025.10.25.684548

**Authors:** Alina Filatova, Olivera Valentirović Dann, Jennifer Bianca Schwarzkopf, Jamina Sofie Gerhardus, Leon Kaysan, Andreas Blaeser, Beate Katharina Straub, Holger Gerhardt, Ulrike A. Nuber

**Affiliations:** Stem Cell and Development Biology, Department of Biology, Technical University of Darmstadt, Germany; Max Delbrück Center for Molecular Medicine in the Helmholtz Association, Berlin, Germany; Institute for BioMedical Printing Technology, Department of Mechanical Engineering, Technical University of Darmstadt, Germany; Centre for Synthetic Biology, Technical University of Darmstadt, Germany; Tissue Biobank of the University Medical Center Mainz, Germany; Institute of Pathology, University Medical Center of the Johannes Gutenberg-University Mainz, Germany; DZHK (German Center for Cardiovascular Research), Berlin, Germany; Charité - Universitätsmedizin Berlin, Berlin, Germany; Berlin Institute of Health, Berlin, Germany; Department of Mechanical Engineering, Technical University of Darmstadt, Germany

**Keywords:** hepatocyte, spheroid, HepaRG, endothelial cells, tissue engineering, HUVEC, serum replacement

## Abstract

Engineering vascularized human liver tissue for *in vitro* models and *in vivo* applications remains a major challenge. Here, we describe a scalable approach to generate human liver spheroids with self-organized, lumen-containing vascular networks and demonstrate their use as building blocks for fabricating vascularized tissue layers. Spheroids were formed from HepaRG liver cells, human umbilical vein endothelial cells (HUVECs), and adipose tissue-derived mesenchymal stem cells (MSCs). The inclusion of MSCs prevented spatial segregation of hepatic and endothelial compartments and enabled endothelial network formation. We present two media that are suitable for culturing these spheroids: a serum-reduced medium and a defined serum-free medium containing Gibco^TM^ KnockOut serum replacement. These media supported the long-term maintenance of hepatocytes in a metabolically active, relatively mature state, as well as the persistence of endothelial networks. Endothelial cell identity and organization were confirmed by VE-cadherin and ICAM-2 immunostaining and by transmission electron microscopy, which revealed adherens junctions and luminal morphologies consistent with a capillary-like organization. Spheroid-derived HUVECs established anastomoses with external endothelial channels in microfluidic devices. Moreover, endothelial sprouts emerging from the spheroids formed inter-spheroid connections within permissive hydrogels (fibrin or collagen–methylcellulose), a process that depended on the inter-spheroid distance. Finally, we demonstrate the fabrication of planar tissue layers with vascularly interconnected spheroids. Together, we identify key conditions, including cellular ratios, medium formulations, biomaterials, and spatial design criteria that enable the generation and assembly of vascularized liver spheroids as scalable tissue building blocks for tissue engineering applications.

**Graphical abstract:** 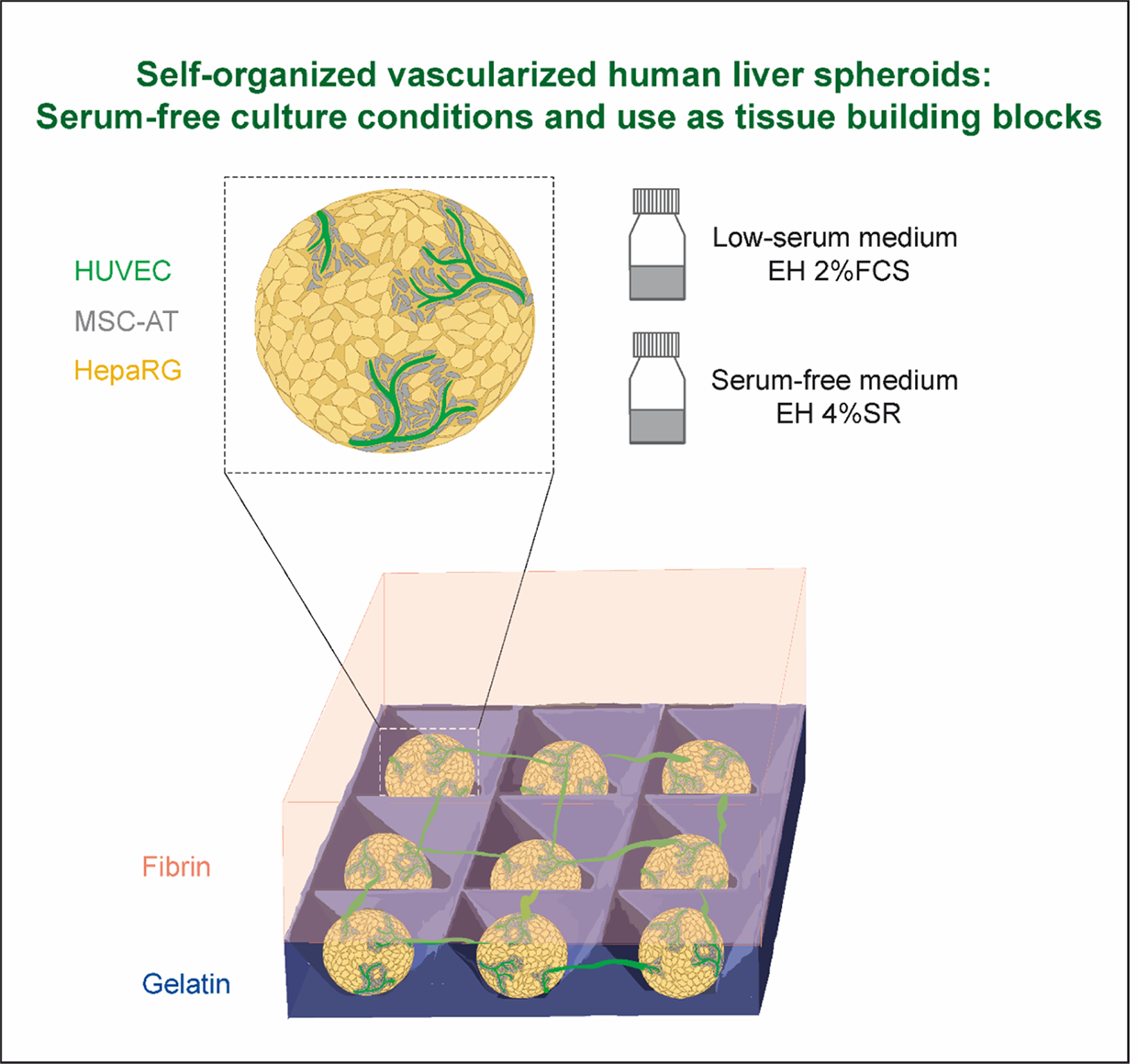

## Introduction

Human tissues generated in the laboratory provide versatile *in vitro* systems for studying human diseases, drug development, and toxicology assessment, and hold great promise for regenerative medicine.^[1, 2]^ They have the potential to reduce animal testing and to serve as an alternative to donated human tissue. Among human tissue types, liver tissue stands out as particularly relevant for drug development, toxicology assessment, and the replacement or compensation of lost or impaired liver function through tissue transplantation. The high medical demand for human liver tissue is also due to critical differences in metabolism and disease characteristics between human and animal liver cells. Drug-induced liver injury is the most frequent cause of the withdrawal of an approved drug, accounting for 30% of all cases, and the insufficient ability of animal models to reflect human liver pathophysiology and drug responses is an important factor for this high failure rate.^[3]^ There is a high medical need for human liver tissue to compensate for or replace dysfunctional one in several severe diseases: chronic liver failure due to chronic viral hepatitis, long-term alcohol abuse, non-alcoholic fatty liver disease, autoimmune hepatitis, and genetic or metabolic disorders. Acute liver failure, often caused by viral hepatitis or toxic agents, as well as inherited liver-based metabolic dysfunction, further contribute to this demand.^[4,5,6]^ However, a routine application of laboratory-generated human liver tissue is hindered by several limitations. One critical deficit of current human tissue models, including those of the liver, is their lack of a vascular system, which severely restricts their utility for various reasons: (i) interactions between vascular and tissue-specific cells are required for normal tissue homeostasis and function and are impaired in various liver diseases, (ii) the vascular system mediates the entry of drugs and pathogens into liver tissue, (iii) a perfusable vascular system is required for tissue models with diameters exceeding the oxygen diffusion limit i.e., of more than several hundred µm. Another key limitation of current human tissue models is their scalability. This is particularly relevant for therapeutic applications of human liver tissue, since sufficiently high tissue mass is needed to compensate for lost function. Bioprinting processes of single cells are currently too slow to fabricate tissue of clinically relevant scale within an acceptable time frame.^[7]^ One appealing strategy to rapidly fabricate larger tissue formations is their assembly from tissue building blocks that already consist of hundreds to thousands of cells.^[8,7]^ For standardized fabrication, such building blocks should ideally have a controllable and uniform size, allow spatially defined deposition and assembly, and be suitable for large-scale production. These requirements are met by spherical three-dimensional (3D) cell cultures (spheroids and organoids) with diameters of a few hundred micrometers. At this size, oxygen and nutrients can still be supplied by the surrounding medium during their production and maintenance. Such 3D cell cultures can principally fuse into larger assemblies.^[9,7]^

The term spheroid, as first described in the 1970s, is mainly used in the literature for spherical 3D cell formations that result from cell aggregation and division.^[10]^ Spheroids can consist of single or multiple cell types. Organoids develop from stem cells through self-organizing processes and therefore usually consist of different derived cell types; the term organoid implies that these 3D cell formations recapitulate specific structural and functional properties of an organ.^[11]^ Many organoid generation protocols produce spherical cell formations, including some that resemble human liver tissue.^[12]^ Of note, the principal applicability of human liver tissue-like spheroids or organoids to restore or replace lost liver functions has been shown upon transplantation into mice.^[13,14,15]^, though much larger constructs would be required for human applications. Since the survival of larger tissue constructs after transplantation depends on rapid vascularization within the tissue, strategies to generate pre-vascularized tissue building blocks are being pursued. Using different approaches, the formation of endothelial cell networks within 3D human liver-like tissue has been shown. However, in very few cases, vascular-like structures that contain a lumen formed by endothelial cells have been achieved, which is a prerequisite for their perfusability (for review, see ^[16]^). In most vascularized organoids or spheroids, luminal structures — i.e., endothelial tubes with a hollow interior — were only observed after transplantation into mice. In these cases, only low numbers of 3D cultures were transplanted and not assembled into larger tissue formations (e.g. ^[17,18]^). Harrison, Siller, Tanaka and Chollet et al. demonstrated endothelial structures with a lumen in liver organoids produced *in vitro* from human pluripotent stem cells (hPSCs) according to light microscopy.^[19]^ Kim and colleagues reported luminal vasculature in liver organoids generated from hPSCs based on *in vitro* live perfusion, light and transmission electron microscopy data.^[20,21]^ Here, we identify various parameters that govern the self-organization of endothelial cells into lumen-containing microvessel-like structures with key hallmarks of natural capillaries in human 3D liver spheroids. We show that human umbilical vein endothelial cells (HUVECs), the most frequently used human endothelial cell type employed in vascularization approaches of 3D cell cultures, do not form a network upon aggregation with human bipotent HepaRG liver cells in spheroids unless an additional mesenchymal cell type is integrated. Furthermore, our experiments demonstrate that defined cell ratios of these three cell types during aggregation into 3D cell clusters allow the formation of vessel-like structures. In addition, we developed novel serum-reduced and serum-free cell culture media that sustain the long-term maintenance of hepatocytes together with HUVEC-formed vascular structures. Moreover, we demonstrate the principal applicability of vascularized liver spheroids as tissue building blocks for the generation of larger tissue constructs, forming endothelial connections between the spheroids and larger endothelialized channels. Finally, we present a methodology for generating tissue layers from evenly distributed spheroids that interconnect via vascular sprouts.

## Methods

### Cell culture experiments

#### Cells and media

All cells were maintained at 37 °C in a humidified 5% CO_2_ incubator. Cryopreserved HepaRG cells (HPR101, Biopredic International, Saint Gregoire, France) were cultured in HepaRG medium (William’s E medium (22551022, Thermo Fisher Scientific, Schwerte, Germany) with 10% fetal calf serum (FCS), 5 µg/ml insulin (I9278, Sigma Aldrich, Darmstadt, Germany), 50 µM hydrocortisone 21-hemisuccinate sodium salt (sc-250130, Santa Cruz Biotechnology, Heidelberg, Germany), 2 mM L-glutamine, 100 U/ml penicillin, and 0.1 mg/ml streptomycin). GFP-labeled HUVEC cells (ZHC-2402, Cellworks, South San Francisco, USA) were maintained in Endothelial Cell Growth Medium 2 (C-22011, PromoCell, Heidelberg, Germany) with 100 U/ml penicillin and 0.1 mg/ml streptomycin. MSC-AT cells (C-12977, Promocell) were grown in Mesenchymal Stem Cell Growth Medium 2 (C-28009, Promocell) with 100 U/ml penicillin and 0.1 mg/ml streptomycin. All cell lines were weekly tested for mycoplasma contamination and were confirmed to be mycoplasma-negative.

#### Generation of master stamps

The master stamps were generated according to Dahlmann et al.^[22]^ by pouring 2.5 ml of silicone (Hydrosil components A and B mixed 1:1, 101301, Siladent Dr. Böhme & Schoeps GmbH, Goslar, Germany) into an AggreWell^TM^800 plate (34811, Stemcell Technologies, Saint-Égréve, France) used as template. Air bubbles were removed from the liquid silicone by centrifugation at 55 x g for 10 seconds using an Eppendorf 5810R centrifuge equipped with a swing-bucket rotor (Eppendorf, Cologne, Germany). The solidified silicone master stamps were gently removed from the AggreWells, washed with cell culture-grade water and 80% ethanol, sterilized by incubation under UV light for 30 min, and air-dried under sterile conditions in a laminar flow hood.

#### Generation of spheroids (s. Fig. 1B)

Pre-heated solution of 2% agarose in William’s E medium was filled into wells of a 12-well plate, and the master stamps were pressed into the agarose. After incubation for 20 min at 4 °C, the stamps were removed, and the plate was sterilized for 30 min under UV light. The cells were seeded into the agarose wells at a density of 1.2 x 10^6^ cells per well in 2 ml of medium with 50 µg/ml gentamicin and incubated for 4 days in the cell culture incubator to form aggregates. Thereafter, the spheroids were transferred in 2 ml of fresh medium into a 6-well suspension plate coated with Anti-Adherence Rinsing Solution (07010, Stemcell Technologies). Medium changes were performed twice a week. Phase-contrast and GFP images of the cultured spheroids were taken using a Zeiss Observer D1 microscope.

In some experiments, the spheroids were cultured in EH 2%FCS medium consisting of 47.75% of Endothelial Cell Basal Medium 2 (C-22211, Promocell), 47.75% of William’s E medium (22551022, Thermo Fisher Scientific), 5 ng/ml EGF (recombinant human), 10 ng/ml basic fibroblast growth factor (bFGF, recombinant human), 20 ng/ml insulin-like growth factor 1 long R3 (IGF-1 LR3, recombinant human), 0.5 ng/ml VEGF165 (recombinant human), 1 µg/ml ascorbic acid, 22.5 μg/ml heparin, 0.2 µg/ml hydrocortisone, 2% FCS (all from the bullet kit C-22111, Promocell), 5 µg/ml insulin 21-hemisuccinate sodium salt (I9278, Sigma Aldrich), 2 mM L-glutamine, 100 U/ml penicillin, and 0.1 mg/ml streptomycin, or EH 4%SR medium containing 46.75% of Endothelial Cell Basal Medium 2 (C-22211, Promocell), 46.75% of William’s E medium (12551, Thermo Fisher Scientific), 5 ng/ml EGF, 10 ng/ml bFGF, 20 ng/ml IGF-1 LR3, 0.5 ng/ml VEGF165, 1 µg/ml ascorbic acid, 22.5 μg/ml heparin, 0.2 µg/ml hydrocortisone (all from the bullet kit C-22111, Promocell), 5 µg/ml insulin 21-hemisuccinate sodium salt (I9278, Sigma Aldrich), 4% Gibco^TM^ KnockOut serum replacement (SR, 10828028, Thermo Fisher Scientific, SR), 2 mM L-glutamine, 100 U/ml penicillin, and 0.1 mg/ml streptomycin.

#### Embedding and culture of vascularized spheroids in microfluidic chips

Triple-cell spheroids were generated five days prior to embedding, according to the protocol above. The spheroids embedded in fibrin gel were seeded into the central chamber of the idenTX3 chip (Aim Biotech, Nucleos, Singapore). RFP-labeled HUVECs (P20216, Innoprot, Bizkaia, Spain) were loaded into one side channel of the chip to form a confluent monolayer acting as parent vessel for the angiogenesis assay. Human brain pericytes (P10363, Innoprot) were seeded into the opposing side channel. The cells were cultured in EH 4%SR medium for 4 days, with 50 ng/ml VEGF165 added to the pericyte-containing side channel to establish a VEGF165 gradient and thereby induce sprouting angiogenesis.

#### Preparation of spheroids for embedding and culture of spheroids in hydrogels

The generation of spheroids was initiated five days prior to embedding according to the protocol above. For embedding, spheroids were washed with calcium- and magnesium-free Dulbecco’s Phosphate-Buffered Saline (dPBS) (D8537-500ML, Sigma-Aldrich) and resuspended in basal medium (BM), composed of a 1:1 mixture of William’s E Medium and Endothelial Cell Basal Medium 2 (C-22011, PromoCell, Heidelberg, Germany). Fifteen to thirty spheroids were transferred into each well of a flat-bottom 96-well plate (96-wp) and kept at 37 °C until gel casting. The spheroids were embedded into hydrogels as specified below and cultured in 200 µL of EH 2%FCS or EH 4%SR medium supplemented with 25 ng/ml human VEGF165 (100-20-2UG, PeproTech^TM^ Thermo Fisher Scientific, Schwerte, Germany). The medium was changed every 1-2 days. Three independent experiments were performed, each with three replicates.

#### Fibrin gel preparation and spheroid embedding

Fibrin gels contained 5 mg/ml fibrinogen (F4883, Sigma-Aldrich), 6 U/ml thrombin (605157-1KU, Sigma-Aldrich) and 42.5% BM. A 15 mg/ml stock solution of fibrinogen (F4883, Sigma-Aldrich) was prepared in 0.9% NaCl (S8776, Sigma-Aldrich), and a stock solution of 40 U/ml thrombin in Tris-buffered saline (PPB001, Sigma-Aldrich). All solutions were thawed immediately before use and kept on ice. The fibrinogen/dPBS mixture was freshly prepared and rapidly mixed with thrombin by pipetting to achieve a final concentration of 10 mg/ml fibrinogen and 10.5 U/ml thrombin (605157-1KU, Sigma-Aldrich). The resulting fibrinogen–thrombin mixture was immediately combined with spheroid-containing BM at a 1.35:1 ratio. The plates were incubated at 37 °C for at least 20 min to allow complete gelation.

#### Preparation of fibrin-agarose–gelatin (FibAgGel) hydrogel and spheroid embedding

FibAgGel blend, which contained 0.35% (w/v) low-melting agarose (A01690, Sigma-Aldrich), 3% (w/v) gelatin (G1890-500G, Sigma-Aldrich), 5 mg/ml fibrinogen, and 5 U/ml thrombin, was prepared by mixing agarose and gelatin (both diluted in dPBS), vortexing the mixture gently and incubating it at 37 °C to equilibrate. Freshly thawed fibrinogen was then added to the agarose–gelatin solution, followed by gentle pipetting to homogenize the hydrogel precursor. For embedding, 72% of the hydrogel precursor was pipetted into each well of a flat-bottom 96-well plate containing 28% of spheroid-containing BM, and gelation was initiated by placing the plate at 4 °C for 10 min. Fibrinogen polymerization was initiated by adding thrombin on top of the gel. The plate was incubated at 37 °C for 20 min to allow complete gelation.

#### Preparation of collagen–methylcellulose (CollMet) hydrogel and spheroid embedding

2.5 mg/ml collagen type I (C3867-1VL, Sigma-Aldrich) was neutralized with ∼9% v/v ice-cold 0.2 N NaOH (1091371000, Merck, Darmstadt, Germany) and 1x Medium 199 (M0650-100ML, Sigma-Aldrich), and mixed at a 1:1 ratio with spheroids in 1.2% methylcellulose (M0512-100G, Sigma-Aldrich, viscosity: 4,000 cP), prepared as described in Tetzlaff et al, 2018^[23]^ in ECGM2 medium (C-22211, PromoCell). All components were maintained on ice prior to mixing to prevent premature gelation. Gels were polymerized by incubation at 37 °C in a humidified incubator with 5% CO₂ for 20– 30 min.

#### Fabrication of gels with evenly distributed spheroids

Negative silicone stamps were made using Aggrewell 800 plates as described above. A 6-mm biopsy punch needle (I4100600, Mediware/Servoprax, Wesel, Germany) was used to cut out a stamp fitting into a well of a 96-well plate. On the day of embedding, a negative silicone stamp was placed onto a thin layer of 5% gelatin in dPBS in a 96-well plate, and the gels were incubated at 4 °C for 15 min. After gelation, transglutaminase (B07B4TYFDJ, Wuerzteufel, Empfingen, Germany) in dPBS was added on top of the gelatin to a final concentration of 4 mg/ml, and the gels were incubated for 15 min at RT to achieve gelatin crosslinking. Spheroids containing 16,000 cells each were loaded into the microwells in a minimal amount of medium. Fibrin gel was then mixed with the spheroids as described above (final volume of 60 µl) and carefully layered over the gelatin matrix, avoiding bubbles and ensuring proper spheroid positioning in the microwells. The plates were incubated at 37 °C for at least 20 min to allow complete gelation. Finally, 200 µl of medium were added on top of the gel, and the medium was exchanged daily.

### Staining of spheroids

Spheroids were fixed in 4% paraformaldehyde (PFA, 0335.3, Carl Roth) in dPBS for 1 h at RT with gentle rotation, washed twice with PBS, dehydrated, and paraffin embedded. The spheroids were cut into 5-µm-thick sections using a Jung RM2055 microtome (Leica Biosystems, Nußloch, Germany) and mounted on Epredia Polysine Adhesion microscope slides (J2800AMNZ, Microm International GmbH, Dreieich, Germany). Prior to stainings, sections were deparaffinized and rehydrated.

*For immunofluorescence staining*, sections were incubated in epitope retrieval citrate buffer (10 mM Sodium Citrate, 0.05% Tween-20, pH 6.0) for 40 min (minutes) in a water bath heated to 100 °C and subsequently cooled for 40-60 min at room temperature (RT). After a washing step in PBS, sections were permeabilized in 0.5% Triton X-100 in PBS for 10 min and blocked with 5% IgG-free BSA (3737.1, Carl Roth, Karlsruhe, Germany) in PBS. Primary antibodies were applied in PBS containing 0.01% Triton X-100 and incubated overnight at 4 °C; PBS with 0.01% Triton X-100 served as negative control. Sections were washed in PBS (3 x 10 min), followed by a one-hour incubation step with respective secondary antibodies diluted in 0.01% Triton X-100 in PBS. Specimens were then washed with PBS (2 x 10 min), counterstained with DAPI (1 μg/ml, 10236276001, Merck, Darmstadt, Germany) for 10 min, washed in PBS (5 min), rinsed in water, dehydrated in ethanol, and air-dried. Mounting was performed with Fluorescence Mounting Medium (S3023, DAKO/Agilent, Waldbronn, Germany). Immunofluorescent images were acquired with Zeiss Observer D1 and Zeiss Axiovert 5 microscopes.

*For immunohistochemical staining*, antigen retrieval was performed in an EDTA-containing buffer (FLEX kit, pH 9, Dako) in a steamer at 100 °C for 30 min. Blocking, staining, and washing were performed in an autostainer (Thermo Scientific) using Dako EnVision FLEX reagents according to the manufacturer’s’ protocols. Sections were mounted with Entellan (1079610500, VWR, Darmstadt, Germany).

*Periodic acid-Schiff (PAS) staining* was performed using standard procedures on a Leica automated stainer. The deparaffinized sections were treated with periodic acid for 4 min, followed by incubation with Schiff’s reagent for 6 min. Subsequently, the slides were washed in sulfur dioxide-containing water for 4 min. Nuclei were counterstained with Gill’s hematoxylin, after which the sections were dehydrated through a graded series of isopropanol, cleared in xylene, and mounted.

*To stain the spheroids embedded in hydrogels*, the gels were washed with PBS (3 x 1 min), and the spheroids in gel were fixed in 4% PFA in PBS for 30 min at RT, followed by three PBS washes. Permeabilization was done in 0.5% Triton X-100 in PBS for 5 min at RT, followed by three washes with PBS. Blocking was done with 3% IgG-free BSA in PBS for 1h at RT. Primary antibodies were applied in PBS containing 3% IgG-free BSA and incubated overnight at 4 °C. Gels were washed three times with PBS, followed by a 1 h incubation with the appropriate secondary antibodies diluted in 3% IgG-free BSA in PBS. Specimens were then washed once with PBS, incubated in 1 μg/ml DAPI for 10 min, washed again with PBS for 5 min, and kept in fresh PBS during imaging. Immunofluorescent images were taken on the same day with a Zeiss Observer D1 microscope.

#### Staining of vascularized spheroids in microfluidic chips

Four days after embedding into the chip, the cells were fixed with 4% PFA for 10 min at RT, followed by washing with 0.5% Triton X-100 in PBS. The samples were blocked with blocking buffer (0.5% Triton X-100, 0.01% sodium deoxycholate, 1% bovine serum albumin (A9418-50G, Sigma-Aldrich), 3% FCS) for 1h at RT. The primary antibodies were diluted in a 1:1 mixture of blocking buffer and PBS and incubated for 3 days at 4 °C with gentle shaking. After washing with PBS containing 0.5% Triton X-100 (3 x 10 min), the chips were incubated overnight with secondary antibodies at 4 °C on a shaker. After washing with PBS containing 0.5% Triton X-100 (3 x 10 min), the chips were incubated with DAPI for 20 min at RT on a shaker, followed by another washing step with PBS. Images were obtained using a 3i spinning-disc confocal microscope equipped with a Zeiss Plan-Apochromat 20x/1.0 NA water-dipping objective and SlideBook imaging software.

### Transmission electron microscopy

Spheroids were fixed in 2.5% glutaraldehyde, stained in osmium tetroxide and embedded in Epon. Ultrathin sections were prepared for microscopy according to routine procedures. Electron microscopy was performed using a Jeol Jem 1400HC microscope (JEOL Ltd., Akishima, Tokyo, Japan) with a mounted 4k camera.

### Image analysis

The projected spheroid area was determined manually using ImageJ (35-69 spheroids per condition). The Ki67-positive and E-cadherin-positive cells were manually counted using the CellCounter plugin of the ImageJ software (https://imagej.net/ij/, National Institutes of Health, Bethesda, MD, USA). To determine the percentages of Ki67-positive and of E-cadherin-positive cells, the number of the counted positive cells was divided by the total number of cells, as determined by DAPI staining. Ten spheroids per condition were analyzed.

The number of spheroids with sprouts in different hydrogels was analyzed in three independent experiments, with three technical replicates each. In addition, the sprout length, defined as the linear distance from the spheroid center to the sprout tip, was measured in three independent experiments, n (fibrin, EH 2%FCS) = 35, n (fibrin, EH 4%SR) = 22, n (FibAgGel, EH 2%FCS) = 8, n (FibAgGel, EH 4%SR) = 6, (CollMet, EH 2%FCS) = 17, n (CollMet, EH 4%SR) = 7. In the setup with defined spheroid positioning, the sprout length was measured for all sprouts emerging from the spheroids placed in nine wells. For 82 spheroid pairs, the inter-spheroid distance, that was defined as the distance between the centers of two spheroids, was measured, and it was noted whether neither spheroid (0/2), only one spheroid (1/2) or both spheroids (2/2) exhibited sprouts. All measurements were performed using ImageJ.

### Statistical analysis

GraphPad Prism 8 (GraphPad software San Diego, CA) was used for statistical tests and graphical display. Normality of data within groups was assessed using the Shapiro–Wilk test. For normally distributed data, group comparisons were performed using the Games-Howell’s multiple comparisons test, which accounts for unequal variances and sample sizes, to determine whether the following datasets were significantly different: projected spheroids area; percentages of Ki-67-positive and E-cadherin-positive cells; percentages of triple-cell spheroids with sprouts cultured in EH 2%FCS or EH 4%SR after embedding in indicated hydrogels; and sprout lengths for the spheroids described above. Significance levels were defined as n.s. (not significant, p > 0.05), *p ≤ 0.05, **p ≤ 0.01, ***p ≤ 0.001. Data are presented as mean ± SEM. Datasets describing whether neither spheroid (0/2), only one spheroid (1/2), or both spheroids (2/2) in spheroid pairs with defined distances exhibited sprouts were not all normally distributed; thus, the Kruskal-Wallis test was applied. Sprout lengths of endothelial cells from positioned triple-cell spheroids cultured in EH 2%FCS or EH 4%SR were compared using an unpaired t-test with Welch’s correction.

## Results

### Mesenchymal cells facilitate endothelial cell network formation within human liver cell spheroids

The HepaRG cell line is a widely used human liver cell line for research, drug development, and toxicology assessment.^[24,25]^ HepaRG cells can be expanded and maintained in culture for extended periods while retaining stable hepatic functions and phenotype, and can differentiate into two distinct liver cell types: hepatocyte-like cells and cholangiocyte-like cells. Endothelial cells such as HUVECs, a well-established and easily accessible primary endothelial cell model from the vein of the human umbilical cord, possess the inherent capacity to self-organize into tube-like structures *in vitro* when placed onto or within extracellular matrix proteins or hydrogel mimics thereof.^[26,27]^

Since HepaRG are principally capable of producing extracellular matrix proteins,^[28]^ we set out to test if HUVECs could form vascular network-like structures in 3D HepaRG spheroids upon aggregation of single cells (Fig. 1). HUVECs stably expressing GFP were used to monitor their position within spheroids by fluorescence microscopy. Single-cell suspensions with HepaRG to GFP-HUVEC cell ratios of 7:2, 5:2, or 5:4 in a medium optimized for endothelial cells, Endothelial Cell Growth Medium-2, and supplemented with 2% FCS (ECGM2 2%FCS), were transferred into microwell plates to generate spheroids containing 4,000 cells each. On day four, formed spheroids were transferred to suspension plates and placed on a shaker (Fig. 1B). In case of all three HepaRG to GFP-HUVEC cell ratios, the two cell types separated over the course of six days, with HUVECs finally located in the spheroid center surrounded by non-fluorescent HepaRG cells (Fig. 1A). Based on these results, we decided to add human mesenchymal stem cells (MSCs) from adipose tissue that are known to produce abundant extracellular matrix, are present in the stromal vascular fraction, and resemble pericytes.^[29]^ These cells are highly accessible and provide a rich and practical source for regenerative medicine and research applications. When MSCs were integrated at a cell ratio of 5:2:2 (HepaRG:GFP-HUVEC:MSC), green fluorescent signals were distributed throughout the spheroid in a network-like pattern until the initial observation period of six days (Fig. 1A).

**Figure 1.**
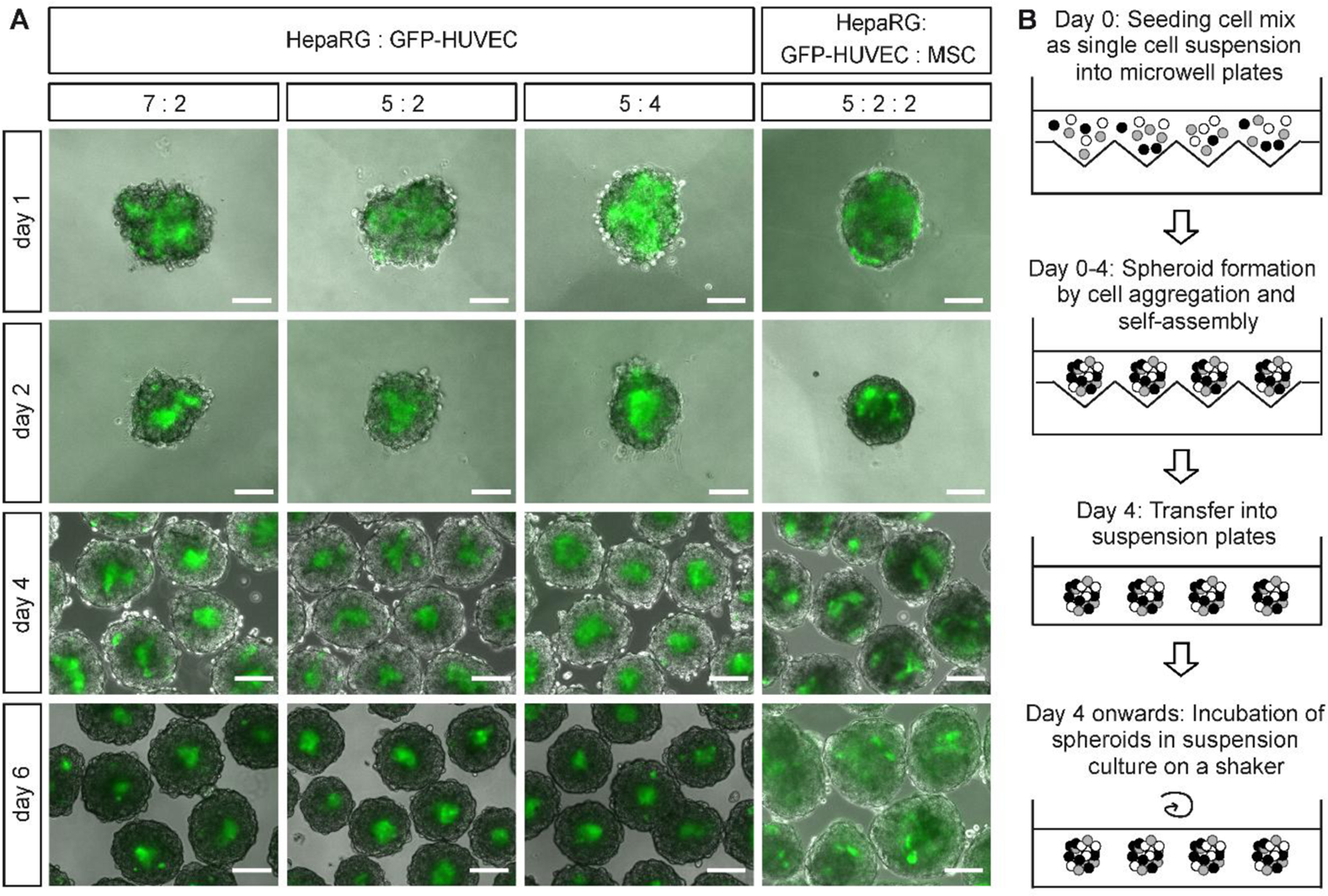
Mesenchymal cells facilitate endothelial cell network formation within human liver cell spheroids. A. Merged bright field and GFP fluorescence microscopy images of spheroids composed of two cell types (HepaRG and GFP-HUVEC) or three cell types (HepaRG, GFP-HUVEC, and MSC) seeded at indicated ratios on days 1, 2, 4, and 6. All spheroids were cultured in ECGM 2%FCS medium. Scale bars: 100 µm. B. Experimental workflow. Mixed single cells were seeded into agarose microwell plates, that were generated using a negative silicone stamp. After four days, the assembled spheroids were transferred into suspension plates and kept on a shaker.

### Vascularized liver spheroid growth is influenced by cell culture medium composition

Although the ECGM2 medium supported the formation of vascular-like structures within the spheroids, the spheroid size decreased over time (Fig. 2A, B, Supplemental Fig. S1), indicating a long-term growth impairment under these conditions. Consistently, these spheroids contained fewer proliferating cells (Fig. 2C, E) and E-cadherin–positive HepaRG cells over time (Fig. 2D, E). To better support HepaRG cells, we adapted the medium composition. In addition to 50% ECGM-2 medium - including growth factors supporting endothelial cells at concentrations equivalent to 100% ECGM2 (5 ng/ml EGF, 10 ng/ml bFGF, 20 ng/ml IGF-1 LR3, 0.5 ng/ml VEGF165, 1 µg/ml ascorbic acid, 22.5 μg/ml heparin, 0.2 µg/ml hydrocortisone) – we added 50% of William’s E medium and 5 µg/ml insulin. This resulting medium was designated EH 2%FCS (Endothelial cell Hepatocyte medium with 2% FCS) to indicate its application for both endothelial cells and hepatocytes. Spheroids cultured in EH 2%FCS were larger (Fig. 2B, Supplemental Fig. S1), contained more Ki67-positive, proliferating cells (Fig. 2C, E), and higher numbers of E-cadherin-positive HepaRG cells (Fig. 2D, E) compared with the spheroids kept in ECGM2 2%FCS. Importantly, the spheroids cultured in EH 2%FCS still retained extensive vascular-like networks (Fig. 2F).

**Figure 2.**
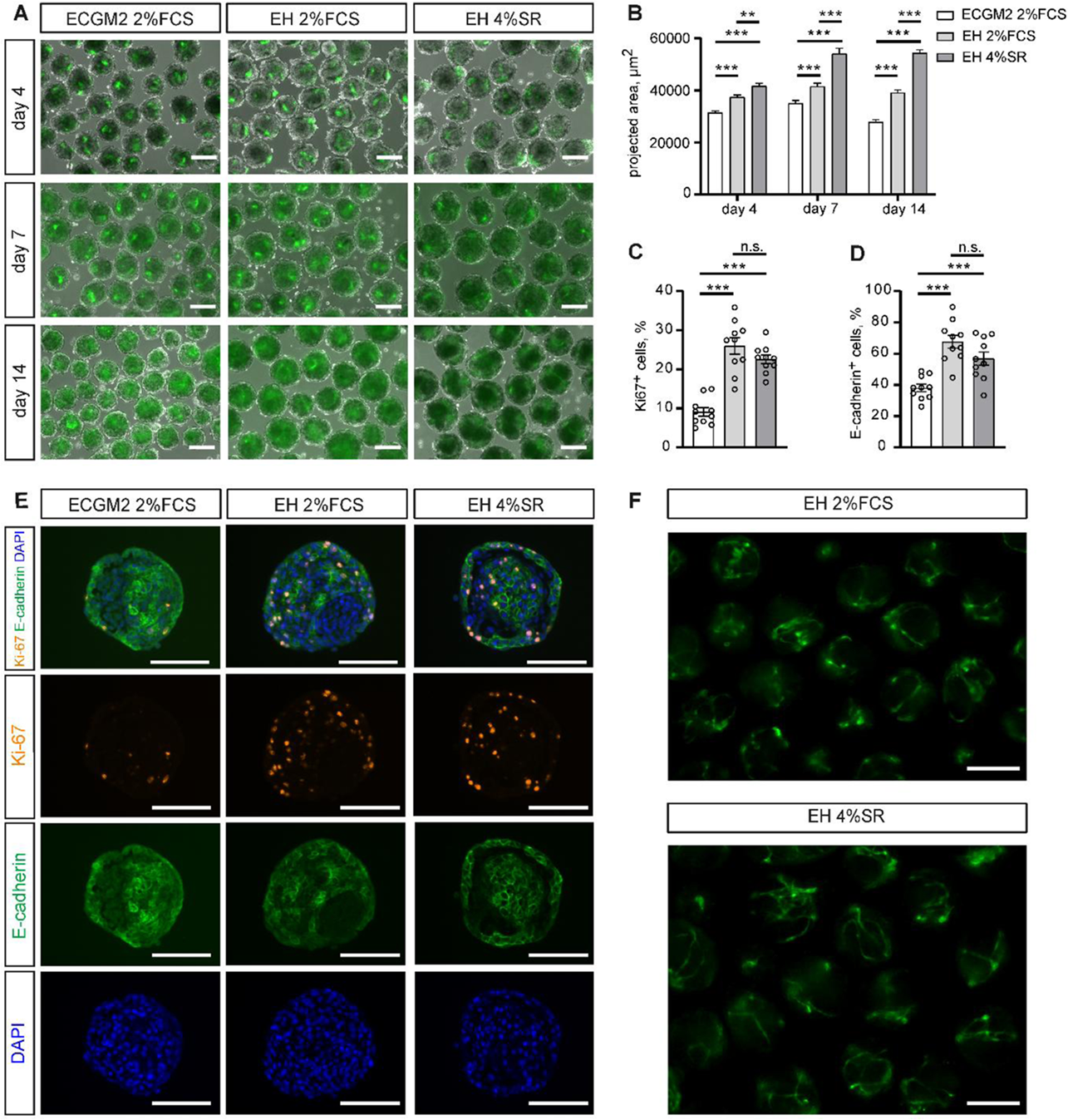
The serum-reduced and serum-free media EH 2%FCS and EH4%SR support triple-cell liver spheroid growth and endothelial cell networks. A. Merged bright field and GFP fluorescence microscopy images of spheroids composed of HepaRG, GFP-HUVEC, and MSC cells at a ratio of 5:2:2 that were cultured in ECGM2 2%FCS, EH 2%FCS, or EH 4%SR for 4, 7, or 14 days. B. Projected area of spheroids cultured in ECGM2 2%FCS, EH 2%FCS, or EH 4%SR on days 4, 7, and 14 after cell seeding. A graphical representation with individual data points is shown in Supplemental Figure S1. (C, D). Numbers of Ki-67-positive (C) and E-cadherin-positive (D) cells within spheroids cultured in ECGM2 2%FCS, EH 2%FCS, or EH 4%SR on day 7 after seeding. E. Representative Ki-67 and E-cadherin immunofluorescence images of paraffin sections of spheroids cultured in ECGM2 2%FCS, EH 2%FCS, or EH 4%SR on day 7 after cell seeding. 4′,6-diamidino-2-phenylindole (DAPI) staining marks the cell nuclei. F. Representative fluorescence microscopy images showing GFP-positive vessel-like structures within spheroids cultured in EH 2%FCS or EH 4%SR for 7 days. Means ± SEM are shown. Scale bars: 100 µm. * p ≤ 0.05, ** p ≤ 0.01, *** p ≤ 0.001, n.s. – non-significant.

We next aimed to identify a serum-free medium with a defined composition that would support the growth of vascularized liver spheroids similarly to the FCS-containing medium. The exclusion of FCS eliminates the batch-to-batch variability inherent in animal serum, reduces the risk of introducing pathogens, aligns with increasing regulatory demands for defined, animal-free media in cell-based therapeutics, and supports the 3Rs principle in research. To this end, we replaced FCS with KnockOut Serum Replacement (SR) in the EH formulation, resulting in EH 4%SR. SR is a chemically defined, serum-free medium supplement widely used to support embryonic stem cell and induced pluripotent stem cell growth without the variability associated with serum.^[30]^ It can also support directed iPSC differentiation and improve survival and yield of desired lineages.^[31,32,33]^ SR contains basic nutrients such as amino acids, vitamins, antioxidants, and trace elements as well as key proteins, namely insulin, transferrin, and lipid-rich albumin. Together, these components support metabolism, redox balance, and growth factor signaling. Insulin and transferrin stimulate cell proliferation and metabolism; albumin is a multifunctional carrier and stabilizer, transporting nutrients and growth factors while protecting cells from toxins and oxidative stress; thiamine, glutathione, and ascorbate support metabolic pathways and protect against stress. We reasoned that these functions would sustain hepatic and endothelial cells *in vitro*. Indeed, the spheroids cultured in EH 4%SR had the largest projected area in comparison to those kept in ECGM2 2%FCS and EH 2%FCS (Fig. 2A, B, Supplemental Fig. S1) and displayed proliferating cells (Fig. 2C, E), E-cadherin-positive HepaRG cells (Fig. 2D, E), and vascular-like structures (Fig. 2F) comparable to the spheroids kept in EH 2%FCS. These results indicate that the replacement of FCS with SR did not impair spheroid growth, cellular proliferation, or maintenance of hepatic and endothelial cell populations. Notably, the presence of vascular-like structures in both conditions suggests that the SR-containing medium supports endothelial organization similarly to the medium with FCS. These findings demonstrate that EH medium supplemented with 4% SR is a viable alternative to traditional FCS-containing media for maintaining the structural and functional integrity of spheroids containing endothelial cells in combination with other cell types.

### Vascularized liver spheroids exhibit characteristics of human liver tissue and can be maintained in culture for three weeks

HepaRG cells are bipotential and can give rise to cholangiocytes and hepatocytes with the latter known to display different stages of maturity under different culture conditions.^(34;35;24)^ To assess the identity and distribution of liver cell types in the triple-cell spheroids and to evaluate long-term cultures, we kept spheroids in EH 2%FCS or EH 4%SR for 14 (Supplemental Fig. S2) or 21 (Fig. 3) days. Immunostaining of paraffin sections from these spheroids revealed expression of a master regulator of hepatocyte differentiation, hepatocyte nuclear factor 4α (HNF4α), along with key hepatocyte maturity markers including cytochrome P450 family 3 subfamily A member 4 (CYP3A4), albumin, and arginase, as well as the presence of proliferating liver cells (Fig. 3A, B; Supplemental Fig. S2A, B). Immunohistochemistry revealed the presence of PAS-positive hepatoid cell trabeculae containing many small lipid droplets (as evidenced by perilipin 2 positivity) in spheroids kept in both media for 21 days, consistent with a metabolically active, relatively mature hepatocytic phenotype (Fig. 3C, D).

**Figure 3.**
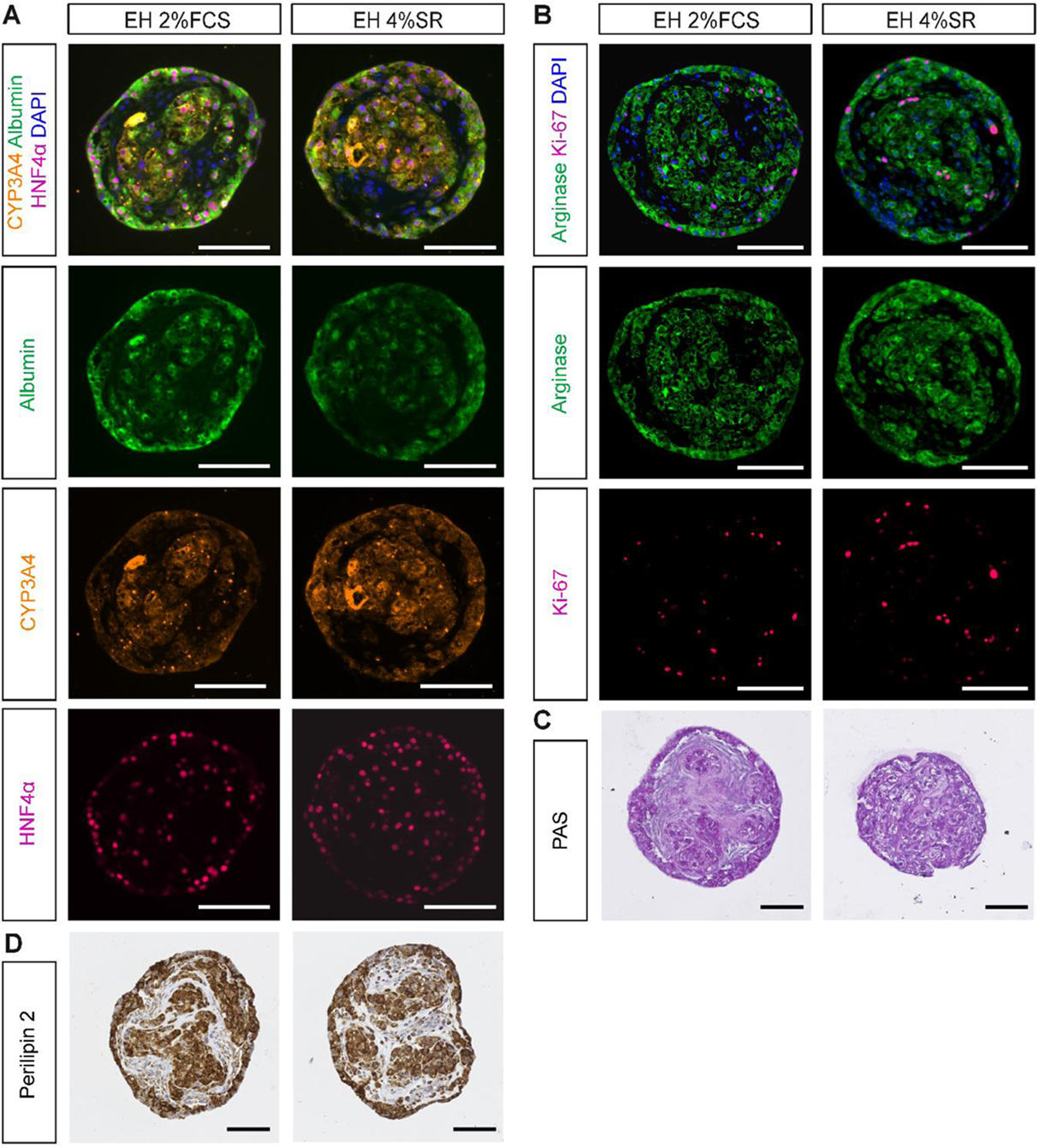
Hepatocyte markers in triple-cell liver spheroids cultured in EH 2%FCS or EH 4%SR for 21 days based on stainings of paraffin sections and PAS reaction. A. Immunostainings with antibodies against albumin (green), CYP3A4 (orange), and HNF4α (magenta). DAPI staining is shown in blue. B. Immunosignals depicting the hepatocyte marker arginase (green), and the proliferation marker Ki-67 (magenta). DAPI staining is shown in blue. C. Periodic acid Schiff (PAS) reaction indicating the presence of glycogen and other carbohydrate-containing entities. D. Immunohistochemistry staining with anti-perilipin 2 antibodies. Scale bars: 100 µm.

Ultrastructural analyses of day 14 (Supplemental Fig. S3A, S4) and day 21 (Fig. 4A, S4) spheroids kept in both media revealed hepatocytic cells with a polygonal shape and well-formed bile canaliculi, typical oval cell nuclei, lipid droplets and glycogen accumulation. In addition, vascular structures embedded in a collagenous matrix were present. The compactness and polygonal shape of hepatocytes on days 14 and 21 were similar; yet, more collagen fibrils were present in EH 4%SR.

**Figure 4.**
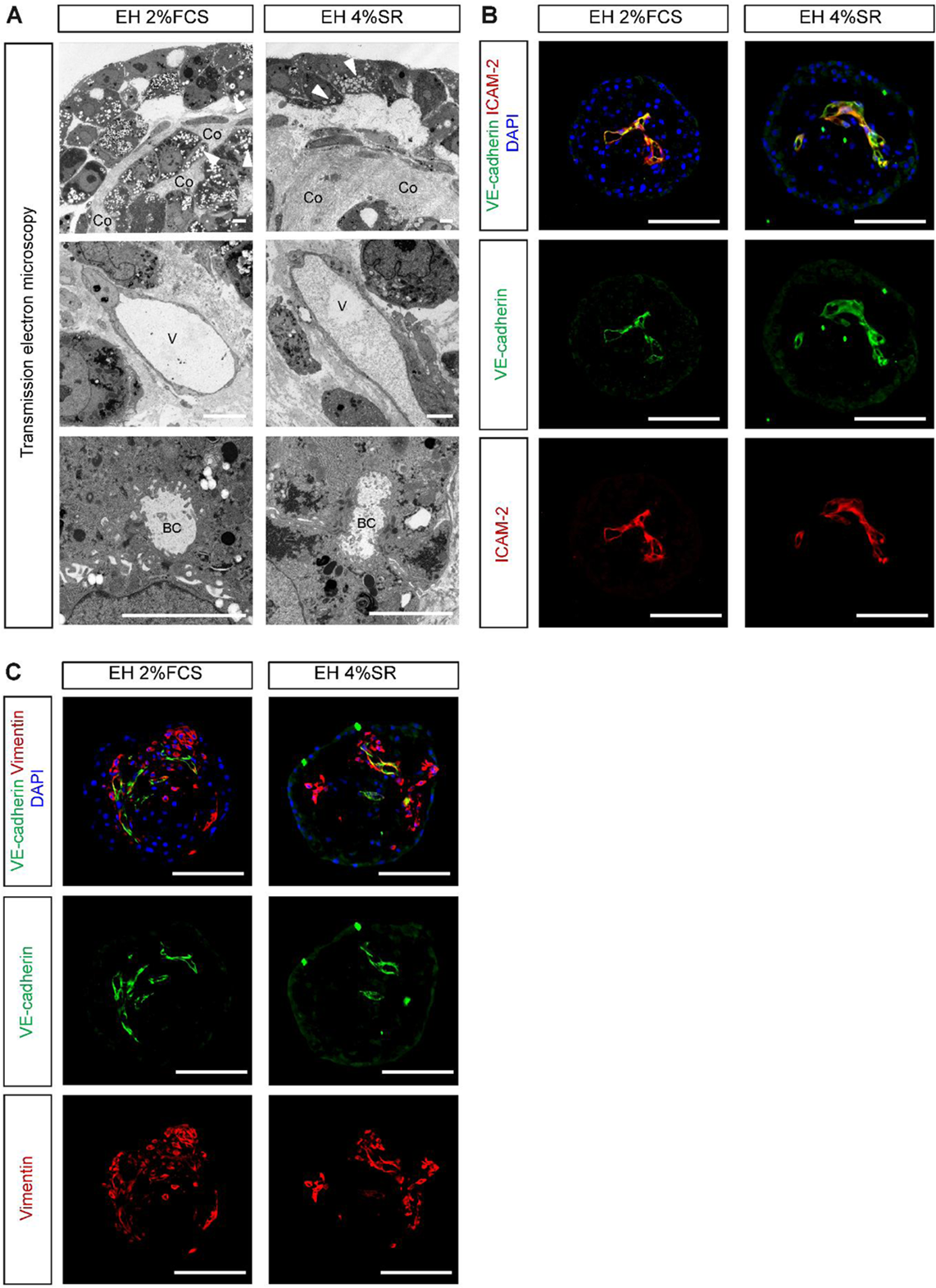
Transmission electron microscopy and immunofluorescence characterization of endothelial structures within the triple-cell liver spheroids. A. Transmission electron microscopy of tissue sections from spheroids cultured in EH 2%FCS or EH 4%SR for 21 days. Upper row: hepatocytic cells at the outer region of spheroids with lipid droplets (white arrowheads) and collagenous stroma (indicated as “Co”). Middle row: capillary-type vessels (V). Bottom row: bile canaliculi (BC) formation in hepatocytic cells. Scale bars: 5 µm. (B, C). Vascular-like structures in spheroids cultured in EH 2%FCS or EH 4%SR for 21 days. Immunosignals of endothelial cell markers VE-cadherin (B, C, green) and ICAM-2 (B, red) and mesenchymal cell marker vimentin (C, red) in spheroid paraffin sections. DAPI staining is shown in blue. Scale bars: 100 µm.

### Endothelial cells form structures resembling microvessels within liver spheroids

HUVECs in the triple-cell liver spheroids formed network-like structures, and lumina within some of these structures were already detectable on day 14 (Supplemental Fig. S3B, S5A) and persisted until day 21 (Fig. 4B) based on VE-cadherin and ICAM-2 immunostainings. The localization of both ICAM-2 and VE-cadherin at HUVEC membranes indicates the establishment of intercellular junctions and a functional endothelial phenotype, supporting cell–cell adhesion, barrier integrity, and vascular-like organization *in vitro*. Transmission electron microscopy of spheroids revealed morphological features that are typical of capillaries: flat endothelial cells interconnected by plaque-bearing adherens junctions in both media on days 14 (Supplemental Fig. S3A, S4) and 21 (Fig. 4A, S4). HUVECs were embedded within the layers of vimentin-positive MSCs, suggesting close cellular interactions between endothelial and mesenchymal cells that may facilitate endothelial integration and support the formation of vascular-like networks within the spheroids (Fig. 4C, Supplemental Fig. S3C, S5B).

### Endothelial cells from liver spheroids form anastomoses with external endothelial cells

To use triple-cell spheroids as building blocks for the generation of larger tissue formations, continuous perfusion through the entire tissue must be achieved. We therefore assessed if the integrated endothelial cells within spheroids can in principle form anastomoses with external endothelial networks by positioning vascularized spheroids next to endothelialized channels in microfluidic chips. GFP-positive HUVECs from spheroids established connections with vessel-like structures formed by RFP-positive HUVECs extending from the surrounding endothelial channel (Fig. 5A). This indicates that the endothelial cells within the spheroid can integrate into external vascular networks, thereby demonstrating their potential to contribute to continuous microvascular structures.

**Figure 5.**
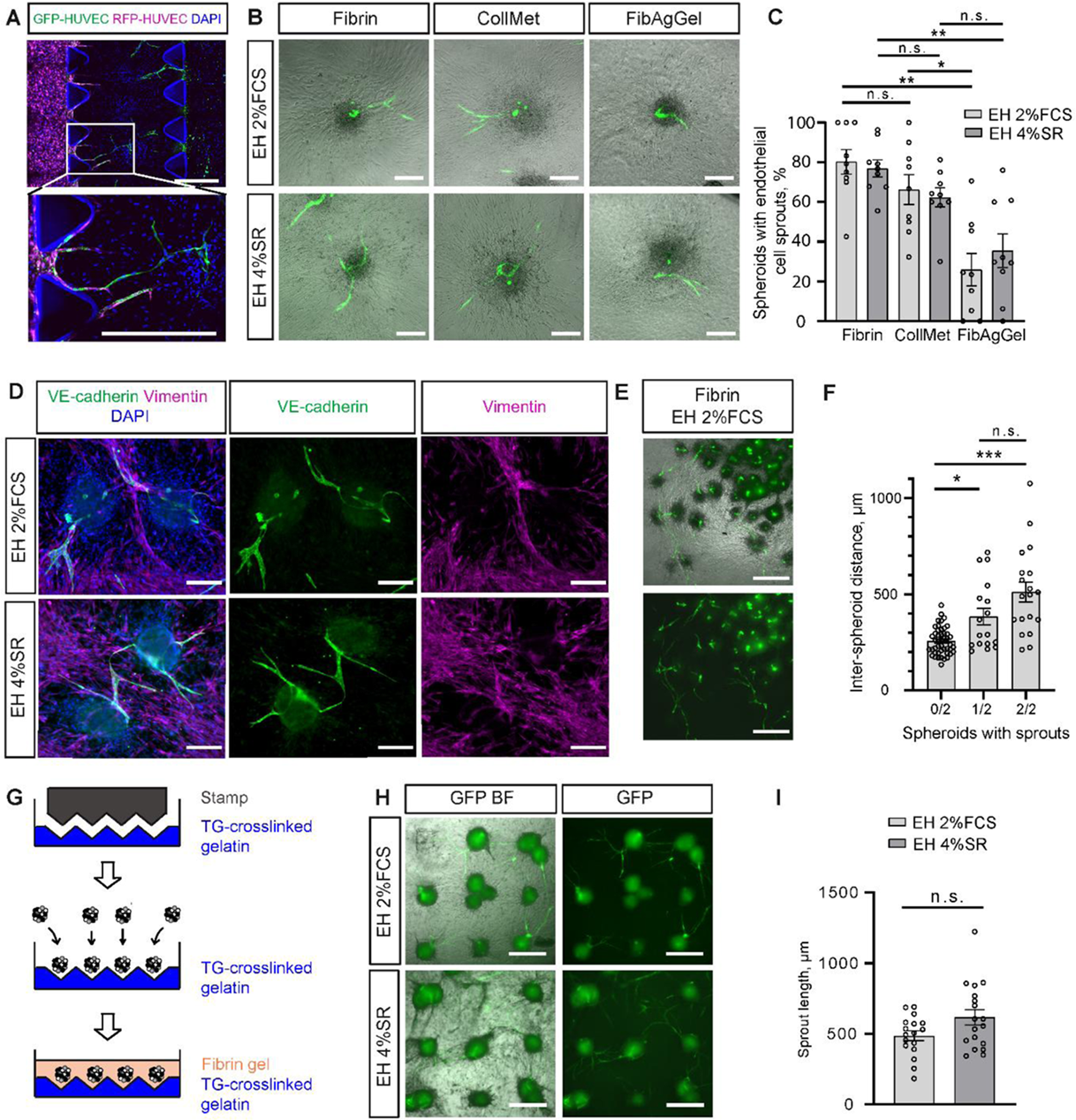
Triple-cell liver spheroid-derived HUVECs formed anastomoses with external endothelial channels in microfluidic devices as well as with HUVECs from adjacent spheroids. A. RFP-expressing HUVECs were loaded into the side channels of a microfluidic chip, and triple-cell spheroids containing GFP-HUVECs were embedded into fibrin in the main chip body. Native RFP signals, signals obtained with anti-GFP antibody and DAPI are shown in magenta, green and blue, respectively. The bottom image represents a magnified area of the labeled area of the upper image. Scale bar: 500 µm. B. GFP fluorescence microscopy images merged with bright field images of triple-cell spheroids cultured in EH 2%FCS or EH 4%SR, embedded in indicated hydrogels on day 2 post embedding. Scale bar: 200 µm. C. Percentage of triple-cell spheroids cultured in EH 2%FCS or EH 4%SR with endothelial cell sprouts on day 3 after embedding in the indicated hydrogels. Means ± SEM are shown; individual data points are indicated. D. Representative immunofluorescence images of triple-cell spheroids embedded in fibrin gel, cultured in EH 2%FCS or EH 4%SR, on day 2 after embedding, and stained for VE-cadherin (green) and vimentin (magenta). DAPI staining is shown in blue. Scale bar: 200 µm. E. Images of GFP fluorescence (bottom) and merged with bright field (top) of triple-cell spheroids cultured in EH 2%FCS on day 4 post embedding into fibrin gel. Scale bar: 500 µm. F. Number of spheroids with endothelial cell sprouts and respective inter-spheroid distance (defined as the distance between the centers of two spheroids). For each spheroid pair, it was noted whether neither spheroid (0/2), only one spheroid (1/2) or both spheroids (2/2) exhibited sprouts. G. Schematic workflow of scalable spheroid embedding with defined inter-spheroid space. A thin gelatin layer with microwells was cast using a negative silicone stamp. After gelation, the gelatin was cross-linked with transglutaminase. Next, spheroids were loaded into the microwells, and fibrin gel was mixed with the spheroids while ensuring their proper positioning in the microwells. H. GFP fluorescence microscopy images (right) and merged with bright field (left), of evenly spaced triple-cell spheroids, cultured in EH 2%FCS or EH 4%SR on day 13 post-embedding into fibrin gel. Scale bar: 500 µm. I. Sprout lengths of endothelial cells from positioned triple-cell spheroids, cultured in EH 2%FCS or EH 4%SR on day 13 post-embedding into fibrin gel. The sprout length was determined as the linear distance from the spheroid center to the end of a sprout. * p ≤ 0.05, ** p ≤ 0.01, *** p ≤ 0.001, n.s. – non-significant.

### Liver spheroids as building blocks for the formation of larger vascularized tissue

We next asked if endothelial cell connections could be formed between multiple adjacent vascularized liver spheroids. This would serve as proof of principle that these spheroids enable a rapid within-tissue vascularization and would underscore their principal applicability as building blocks to assemble larger tissue formations. For this purpose, endothelial cell sprouting between the spheroids had to be facilitated. Initial experiments with spheroids randomly placed in different types of hydrogels yielded several important results. First, among the three hydrogels tested (fibrin, collagen-methylcellulose (CollMet), fibrin-agarose-gelatin blend (FibAgGel)), fibrin and CollMet gels proved most effective, yielding the highest number of spheroids with sprouts (Fig. 5B, C). There was a trend towards longer maximal sprout length in fibrin and CollMet gels compared with FibAgGel, reaching statistically significant differences in case of fibrin vs FibAgGel in medium EH 2%FCS and CollMet vs FibAgGel in EH 4%SR (Supplemental Fig. S6). Second, we observed that the migration of spindle-shaped GFP-negative cells out of the spheroid often preceded the appearance of GFP-positive endothelial cell sprouts. Immunostaining of the fibrin gel-embedded spheroids revealed vimentin-positive MSCs in the areas surrounding the spheroids, while VE-cadherin-positive endothelial cells extended towards these MSCs (Fig. 5D). Third, endothelial sprouts occurred mainly at spheroid sites where the distance to neighboring spheroids was larger, i.e. no sprouts were detected between spheroids placed less than 200 µm apart, and the permissive distance lied between 205 and 1075 µm (Fig. 5E, F).

In final experiments, the inter-spheroid sprouting of endothelial cells was studied and exploited to generate a tissue layer composed of multiple vascularly interconnected spheroids. Based on the previously derived results (Fig. 5B, C, E, F), we defined design criteria to fabricate one tissue layer assembled from multiple vascularized liver spheroids interconnected by endothelial cells: spheroids should be evenly spaced, with a distance of 200-1000 µm between neighboring spheroids, and bridges of either fibrin or CollMet gel should facilitate endothelial cell sprouting between them. To ensure even spacing between spheroids, we fabricated a negative silicone stamp to cast a thin layer of gelatin with evenly distributed wells for spheroid placement (Fig. 5G), leading to an inter-spheroid distance of 839 ± 34 µm horizontally and vertically, and 982 ± 20 µm diagonally (both means ± SEM). Fibrin gel containing spheroids was pipetted onto the gelatin wells, ensuring an even distribution of 1–3 spheroids per well and the formation of fibrin bridges between the spheroid-containing wells (Fig. 5G). In this setup, the gelatin wells served two functions: (1) They could be readily cast with defined size and distance, which was essential for the controlled seeding of spheroids at the beginning of the experiment. (2) During cultivation of the spheroids, the gelatin base mechanically reinforced the surrounding fibrin matrix. To maintain structural support, the gelatin matrix was cross-linked using transglutaminase. This approach ensured that spheroids remained confined within their designated wells and maintained defined inter-spheroid spacing throughout the culture period. Without transglutaminase treatment, the patterned gelatin transitioned from the gel to the sol state and dissolved under cell culture conditions. As a result, shrinkage of the remaining fibrin gel could be observed, which – without the previous mechanical support through the gelatin matrix – could not withstand the contractile forces of the embedded cells, a frequently reported challenge using this protein-based hydrogel in high-density 3D cell cultures. The shrinkage occurring in the absence of trans-glutaminase treatment altered the well geometry, led to irregular spheroid positioning, and caused a loss of defined spheroid spacing (Supplemental Fig. S7), thereby compromising the reproducibility of the experimental setup. The methodology described was then applied to study the formation of endothelial sprouts between spheroids under controlled spatiotemporal conditions. The first sprouts appeared 5-7 days after embedding and vascular inter-connections had formed 13 days post-embedding (Fig. 5H, I), as evidenced by the fusion of GFP-HUVEC sprouts from neighboring spheroids. The endothelial sprouts had a length of 511 ± 35 µm and 594 ± 55 µm (both means ± SEM) in EH 2%FCS and EH 4%SR, respectively, with no significant differences observed between the two studied media compositions (Fig. 5I). These results prove the general feasibility of the proposed concept for the use of the triple-cell spheroids as building blocks for the formation of larger vascularized tissue layers.

## Discussion

Vascularized human liver spheroids hold great potential in the hepatology field: as tissue models to study liver diseases, assess substance toxicology, and develop drugs, as well as tissue building blocks to rapidly fabricate larger tissue units that can replace or support lost or dysfunctional human liver tissue. Our study identifies key parameters that influence the coordination of multiple self-organizing processes that occur simultaneously in space and time upon the aggregation of different cell types into spheroids, ultimately resulting in the formation of vascularized human liver tissue. We furthermore present a new methodology that enables the formation of larger, contiguous, vascularized tissue formations from spheroids by creating conditions for inter-spheroid vascular connections.

Our results outline the key requirements for establishing long-term cultures of self-assembled vascularized liver spheroids. (i) The inclusion of a mesenchymal cell type into spheroids prevents individual liver and endothelial cells from forming two spatially separated, aggregated cell groups, with central and peripheral spheroid localization, respectively, and enables the endothelial cells to organize themselves into network-like structures. (ii) A numerical ratio of these three cell types (5:2:2 of HepaRG:GFP-HUVEC:MSC) leads to consistent vascularized spheroid formation. (iii) An exquisite challenge of maintaining spheroids consisting of cells that are typically grown in very different media formulations is to identify a common medium that accommodates the requirements of all included cell types. For liver spheroids based on HepaRG cells and HUVECs, we show that under the described serum-reduced and serum-free culture conditions, the HepaRG-derived hepatocytes exhibit a metabolically active, relatively mature phenotype according to hepatocyte marker expression and transmission electron microscopy. (iv) Moreover, these culture conditions support the development of endothelial cell networks with the potential of lumen formation. Importantly, we identified endothelial cell formations with ultrastructural features of capillaries. This result is a notable achievement since for the majority of reported liver spheroids, lumina only form upon transplantation,^[16–18]^ and of the ones describing lumina, we are only aware of one group that has provided ultrastructural images of formed vessels.^[20,21]^ In contrast to our work, the cells used in these two studies were derived from human embryonal and human induced pluripotent stem cells.

Based on our results, we suggest that the described approach could have broad applicability and may serve as a template for generating other human spheroids consisting of an organ-specific cell type, endothelial cells, and a mesenchymal cell type, namely to apply a 1:1 mixture of basal media used for endothelial cells and the organ-specific cell type supplemented with 100% final concentrations of all additives required for the two cell types, with a potential use of KnockOut serum replacement instead of FCS. Importantly, our spheroid generation and maintenance protocol does not necessitate any additional extracellular matrix or hydrogel and is easily scalable.

Most studies on spheroids as tissue building blocks have focused on fabrication methods that promote a narrow spheroid arrangement, facilitating fusion between neighboring spheroids. However, in case of vascularized spheroids, such short inter-spheroid distance might be disadvantageous. We demonstrate that the distance between spheroids is a critical factor for the formation of endothelial connections between vascularized liver spheroids, and that spheroids placed in close proximity fail to establish vascular interconnections. These results are consistent with the findings of Kim et al.^[35]^ who demonstrated that a distance of 200 µm between spheroids generated from MSCs derived from human adipose tissue, alone or in combination with HUVECs, was required for MSC migration into the surrounding Matrigel and for the formation of cellular bridges to neighboring spheroids.^[35]^ Similar to Kim et al,^[35]^ we show that MSCs appear to act as guiding cues for endothelial outgrowth, supporting their directed outgrowth toward adjacent spheroids. These observations challenge the prevailing aim to achieve larger tissue formations by tightly packing spheroids as tissue building blocks. Instead, they underscore the need for automated technologies that allow precise positioning of spheroids with defined inter-spheroid distance. This approach could provide an additional advantage, namely potential functionalizations of the inter-spheroid space, for example by incorporating biomaterials or supportive cell types to further promote vascular integration and tissue function. In addition, spheroid placement strategies, including the microneedles-based (“Kenzan”) spheroid assembly method,^36^ could benefit from the acquired insights on spheroid spacing to facilitate inter-spheroid vascular connections. Irrespective of the fabrication method, creating spaces between adjacent spheroids reduces the required spheroid number, and using pre-fabricated spheroid-containing gel layers can accelerate the fabrication process. Ultimately, leaving space between spheroids as modular building blocks should allow the construction of tissues with cell densities approaching those of native organs, since spheroids inherently contain a high cellular density.

We provide a methodology that allows the production of a tissue layer of vascularly connected liver spheroids, which involves simply seeding prefabricated spheroids within a fibrin gel into a hydrogel layer with cavities, that are quickly generated using a stamp. The resulting plane fibrin surface makes this method potentially amenable to simple, automated, and rapid layer-by-layer fabrication. In addition, we present the need for suitable hydrogels that enable a vascular interconnection of adjacent spheroids through endothelial cell sprouting and identify fibrin and collagen-methylcellulose as suitable ones. Combining self-organization of multiple cell types into vascularized liver spheroids with this methodology may enhance the *in vitro* survival of engineered tissue constructs and may open new ways to scale up engineered tissues. Future investigations should explore the inclusion and maintenance of additional cell types, such as immune cells, either within spheroids or in the inter-spheroidal space, according to the protocols and cell culture conditions described here. Moreover, the *in vitro* perfusability of tissue constructs engineered in this way should be assessed. Furthermore, the potential for layer-by-layer fabrication of macroscale tissue using these vascularized spheroids and our described methodology should be explored. For *in vivo* applications, that depend on rapid vascularization upon implantation, our methodology of building vascularly interconnected tissue blocks may improve the survival of transplanted tissue built from spheroids or organoids.

## List of abbreviations

3D: three-dimensional
AFP: Alpha-1-Fetoprotein
bFGF: basic fibroblast growth factor
BM: basal medium
CollMet: collagen–methylcellulose
CYP3A4: cytochrome P450 family 3 subfamily A member 4
dPBS: Dulbecco’s Phosphate-Buffered Saline
EH 2%FCS medium: Endothelial cell Hepatocyte medium with 2% FCS
EH 4%SR medium: Endothelial cell Hepatocyte medium with 4% KnockOut serum replacement
FibAgGel: fibrin-agarose–gelatin
FCS: fetal calf serum
HSCs: hepatic stellate cells
HNF4α: hepatocyte nuclear factor 4α
h: hour
hPSCs: human pluripotent stem cells
IGF-1 LR3: insulin-like growth factor 1 long R3
HUVECs: human umbilical vein endothelial cells
min: minute
MSCs: mesenchymal stem cells
PAS staining: periodic acid-Schiff staining
PFA: paraformaldehyde
RT: room temperature
SMA: smooth muscle actin
SR: KnockOut serum replacement

## Acknowledgements

Assistance with the study. We are very grateful to Philip Hewitt (early Investigative Toxicology, Merck KGaA), Thomas Herget (Silicon Valley, Innovation Hub, Merck KGaA), and Michael Potente (Max-Delbrück-Center and Charité, Berlin) for discussion and advice.

Meike Stotz-Reimers, Liliana Davkova, and Michaela Becker-Roeck (Stem Cell and Development Biology, Technical University of Darmstadt) as well as Karin Molter (electron microscopy facility, University Medical Center Mainz) and Bonny Adamy (Tissue Biobank, University Medical Center Mainz) are highly acknowledged for excellent technical support.

Presentation: none.

