## Supplemental data for "Self-organized vascularized human liver spheroids: Serum-free culture conditions and use as tissue building blocks"

**Table S1.** Primary antibodies and conditions used in immunostainings.

| Primary antibody | Source | Dilution |
|---|---|---|
| Albumin | A80-229F, Bethyl (Montgomery, Texas, USA) | 1:200 |
| Arginase | 66129-1-Ig, Proteintech (Martinsried, Germany) | 1:200 |
| CYP3A4 | MA5-17064, Thermo Fisher Scientific | 1:100 |
| E-cadherin | AF648, R&D Systems | 1:400 |
| GFP | ab13970, Abcam | 1:300 |
| HNF4α | 3113, Cell Signaling Technology (Leiden, The Netherlands) | 1:200 |
| ICAM-2 | 13355, Cell Signaling Technology | 1:200 |
| Ki67 | ab16667, Abcam (Waltham, Massachusetts, USA) | 1:100 |
| Perilipin 2 | 610102, Progen Biotechnik (Heidelberg, Germany) | undiluted |
| Smooth muscle actin | IR61161-2, Agilent | undiluted |
| VE-cadherin | AF938, R&D Systems (Minneapolis, Minnesota, USA) | 1:100 |
| Vimentin | 5741, Cell Signaling Technology | 1:200 |

**Table S2.** Secondary antibodies and conditions used in immunostainings.

| Secondary antibody | Source | Dilution |
|---|---|---|
| donkey anti-goat-AF488 | 705-545-147, Dianova | 1:300 |
| donkey anti-mouse-AF647 | 715-605-151, Dianova | 1:300 |
| donkey anti-mouse-Cy3 | 715-165-150, Dianova | 1:300 |
| donkey anti-rabbit-AF647 | 711-605-152, Dianova | 1:300 |
| donkey anti-rabbit-Cy3 | 711-165-152, Dianova | 1:300 |
| donkey anti-chicken-AF488 | 703-545-155, Dianova | 1:300 |

### Supplemental figures

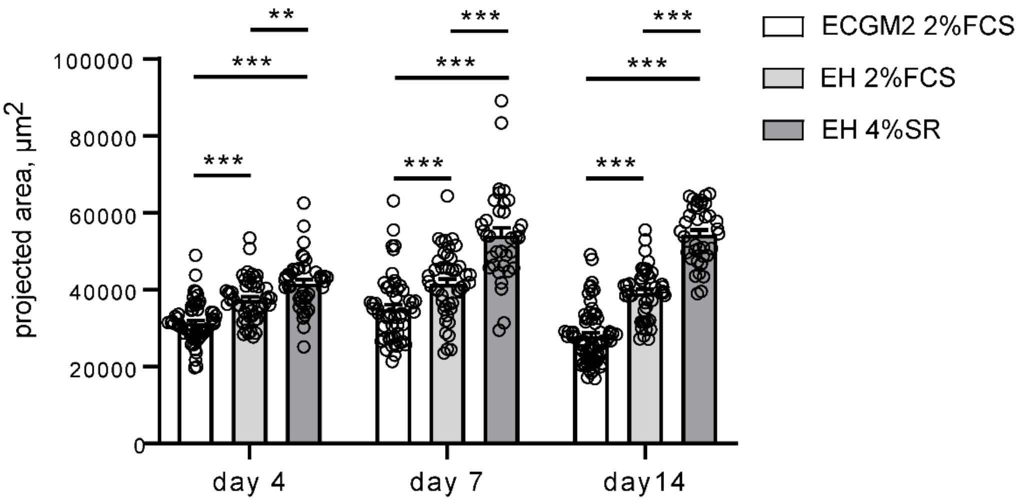


**Figure S1.** Graphical representation of Figure 2B, albeit with individual data points shown. Projected area of spheroids cultured in ECGM2 2%FCS, EH 2%FCS, or EH 4%SR on days 4, 7, and 14 after cell seeding. Means ± SEM are shown. ** $p \leq 0.01$, *** $p \leq 0.001$.

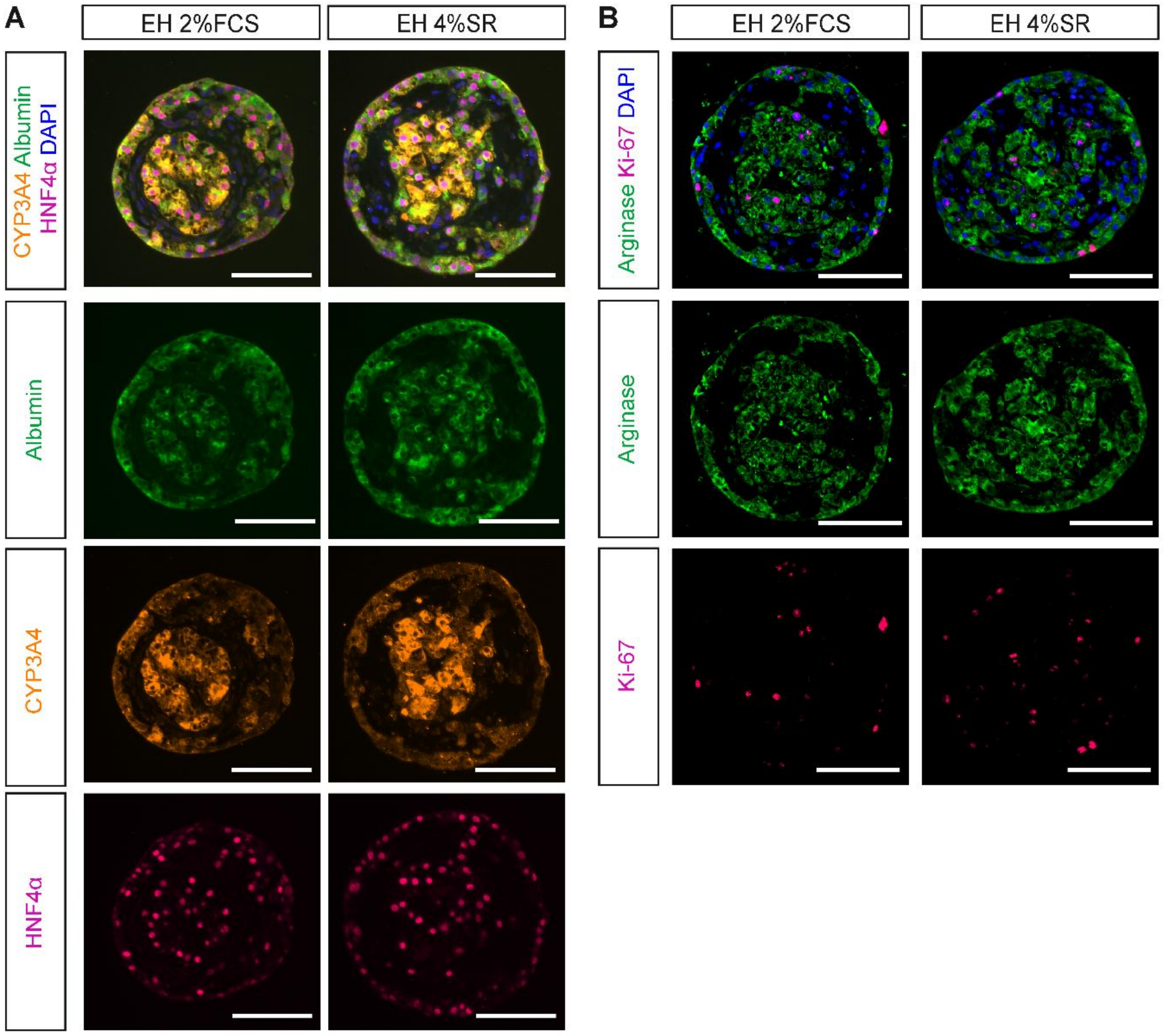


**Figure S2.** Hepatocyte markers in triple-cell liver spheroids cultured in EH 2%FCS or EH 4%SR for 14 days based on stainings of paraffin sections. A. Immunostainings with antibodies against albumin (green), CYP3A4 (orange), and HNF4α (magenta). DAPI staining is shown in blue. B. Immunosignals depicting the hepatocyte marker arginase (green) and the proliferation marker Ki-67 (magenta). DAPI staining is shown in blue. Scale bars: 100 μm.

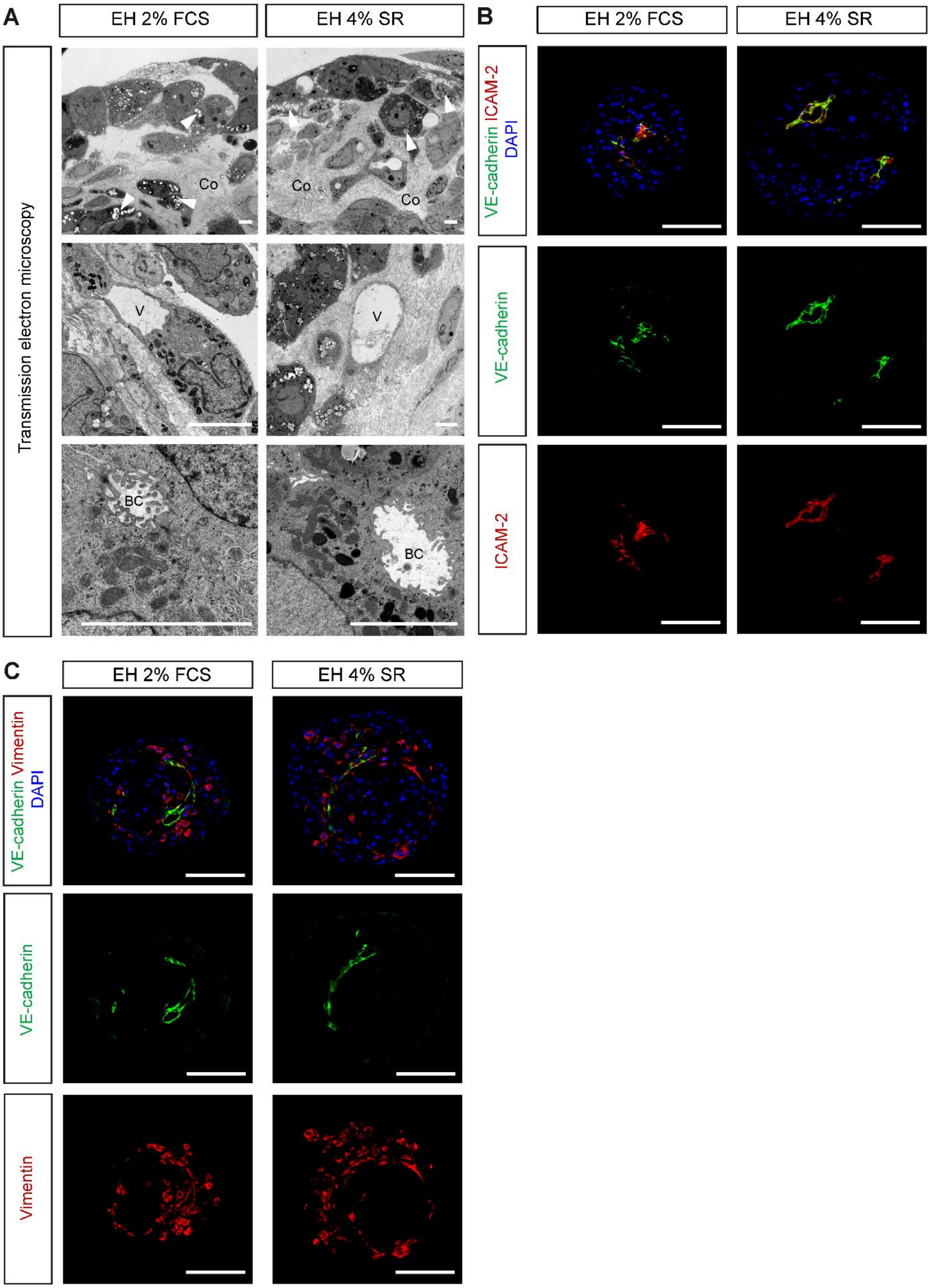


**Figure S3.** Transmission electron microscopy and immunofluorescence characterization of the endothelial structures within triple-cell liver spheroids. A. Transmission electron microscopy of tissue sections from spheroids cultured in EH 2%FCS or EH 4%SR for 14 days. Upper row:

hepatocytic cells at the outer region of spheroids with lipid droplets (white arrowheads) and collagenous stroma (indicated as “Co”). Middle row: capillary-type vessels (V). Bottom row: bile canaliculi (BC) formation in hepatocytic cells. Scale bars: 5 µm. (B,C). Vascular-like structures in spheroids cultured in EH 2%FCS or EH 4%SR for 14 days. Immunosignals of endothelial cell markers VE-cadherin (B, C, green) and ICAM-2 (B, red) and mesenchymal cell marker vimentin (C, red) in spheroid paraffin sections. DAPI staining is shown in blue. Scale bars: 100 µm.

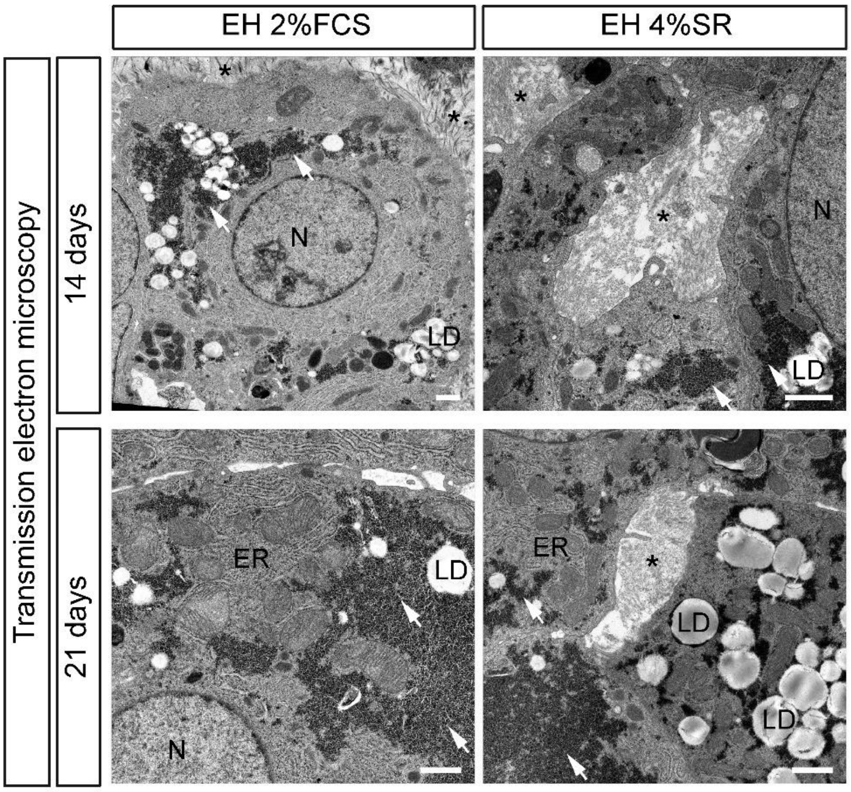


**Figure S4.** High-magnification transmission electron microscopy images of tissue sections from triple-cell liver spheroids cultured in EH 2%FCS or EH 4%SR for 14 or 21 days. The following structures are shown: cell nucleus (N), glycogen (white arrows), lipid droplets (LD), rough endoplasmic reticulum (ER), collagenous stroma (asterisk). Scale bars: 1 µm.

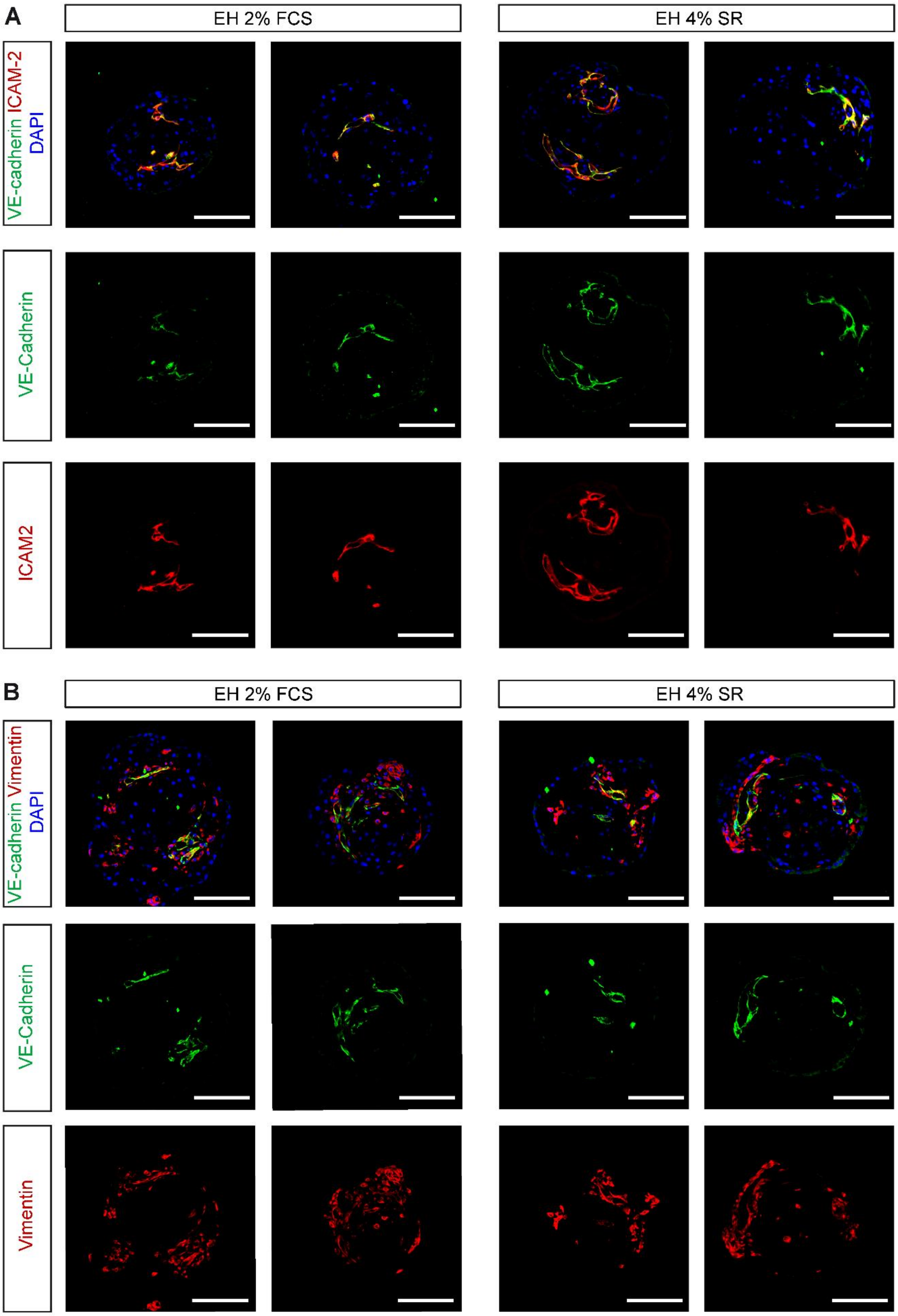


**Figure S5.** Vascular-like structures in triple-cell liver spheroids cultured in EH 2%FCS or EH 4%SR for 21 days. Immunostainings of paraffin sections from spheroids with antibodies against endothelial cell markers VE-cadherin (A, B, green) and ICAM-2 (A, red) and mesenchymal cell marker vimentin (B, red). Scale bars: 100 µm.

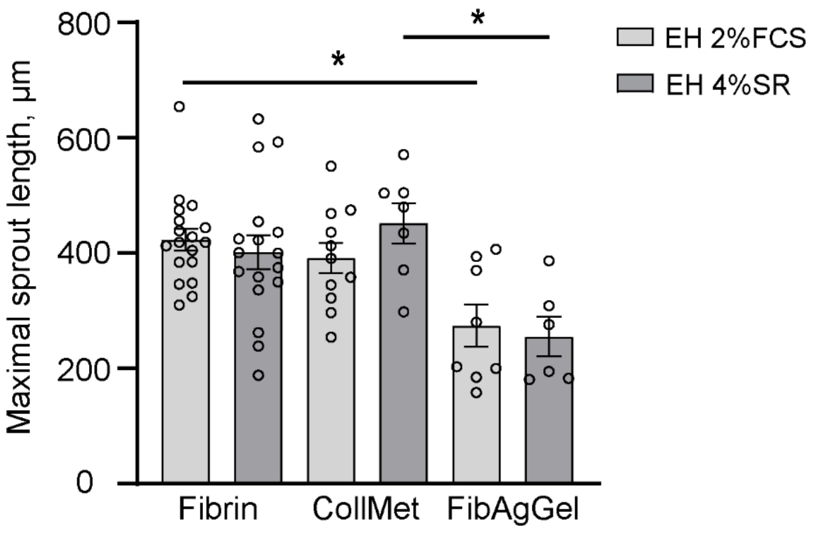


**Figure S6.** Maximal sprout length in triple-cell liver spheroids cultured in EH 2%FCS or EH 4%SR on day 3 after embedding in the indicated hydrogels. Sprout length was determined as the linear distance from the spheroid center to the end of a sprout. Means ± SEM are shown; individual data points are indicated. Significantly different values for each medium are marked; non-significant differences are not indicated. * $p \leq 0.05$.

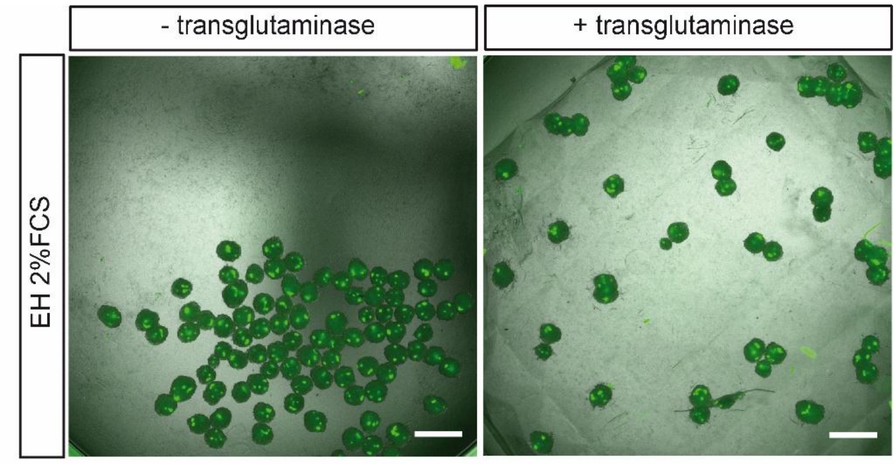


**Figure S7.** In the absence of trans-glutaminase treatment, gelatine shrinkage leads to an altered well geometry and a loss of defined spheroid spacing. Merged bright field and GFP fluorescence images of triple-cell liver spheroids cultured in EH 2%FCS right after embedding into fibrin gel and incubation at 37°C, without (left) and with (right) transglutaminase added to crosslink gelatin microwells. Scale bar: 500 µm.
